# MDA-MB-231 cell morphology influences chemotactic sensing of CXCL12 gradients in type 1 bovine collagen matrix

**DOI:** 10.64898/2026.02.04.703810

**Authors:** Marsophia Marcellus, Catherine J. Murphy

## Abstract

Chemotaxis plays a critical role in the metastatic progression of breast cancer. The chemokine CXCL12 is well recognized as an essential component of chemotactic migration in triple-negative breast cancer (TNBC) cells *in vivo*. The purpose of this study is to determine how the highly metastatic TNBC cell line, MDA-MB-231, migrates in response to well-defined CXCL12 gradients *in vitro*. Traditional 2D transwell migration assays were optimized to gauge the MDA-MB-231 cells’ responsiveness to various CXCL12 concentrations. The optimum chemoattractant concentrations were applied to a commercially available 3D chemotaxis assay as stable linearly diffused gradients. Cells were embedded in type 1 bovine collagen at two different collagen concentrations, and individual unlabeled cells were monitored for 24 hours using brightfield microscopy. Time-lapse videos were used to track cell movement and shape. Quantitative data analysis was performed using an automated tracking software to measure chemotactic parameters based on cell morphology. MDA-MB-231 cells were responsive to CXCL12 concentrations greater than 200 ng/mL in 2D and 3D systems. In 3D systems, significant directed migration was observed in denser collagen matrices. It was observed that in 3D matrices a range of cell morphologies was present. Therefore, chemotaxis was evaluated as a function of cell shape revealing some differences between sub cellular populations. Our findings show the cell’s shape influences the chemotactic sensing towards CXCL12 gradients.

## Introduction

Chemotaxis, the directed migration of cells in response to a chemical stimulus, is responsible for many physiological events such as angiogenesis, wound healing, and cancer metastasis [1]. Chemotactic migration of cancer cells is initiated by the release of chemoattractants, such as chemokines and growth factors, from neighboring cells within the extracellular matrix (ECM). At the onset of metastasis, the associated cell surface receptors to the chemoattractants assist the invading primary tumor cells to sense soluble chemical gradients, to aid in directed migration into the bloodstream and proliferation at a secondary site [1,2] For triple-negative breast cancer (TNBC) cells, where there is a lack of estrogen, progesterone, and human epidermal growth factor 2 receptors, there are a limited number of chemoattractants that could promote their migration. Previous studies have shown epidermal growth factor receptor (EGFR) and G-protein coupled receptor, CXCR4, are prevalent in the TNBC cell line, MDA-MB-231 [3–6]. CXCR4, which has only one known associated ligand, CXCL12, has been extensively studied using *in vitro* migration assays [7–10].

Transwell migration assays, a two-chamber system separated by a porous membrane, have been widely used to measure chemotaxis for two-dimensional (2D) migration. Several studies have conducted CXCL12-mediated chemotaxis of MDA-MB-231 cells with this assay in a high-throughput screening manner [3,7,11–13]. However, these previous findings have reported various cell densities, concentration ranges, and serum exposure times, making it challenging for comparative analysis [14] Although 2D migration assays have brought fundamental insight into TNBC cell migration, it does not reflect how these cells behave in a three-dimensional (3D) ECM environment [15]. Modern-day *in vitro* chemotaxis assays have adapted to 3D cell culture systems, where TNBC cells are embedded in a 3D matrix, primarily made of type 1 collagen, and exposed to a linearly diffused gradient [16–19]. In 2013, Kim et al. discovered MDA-MB-231 cells displayed chemotactic migration when exposed to CXCL12 gradients in a microfluidic system containing type 1 rat tail collagen, suggesting that CXCL12 alone does promote directed migration [20]. However, since then, new reports have shown that the physical properties of the collagen matrix play a vital role in cell migration.

Collagen stiffness, pore size, and source material aid in the cells’ ability to migrate in a 3D system [21]. Type 1 bovine collagen is an alternative material used for 3D cell culture [22]. Fibril morphology studies using confocal microscopy found the organization of collagen fibrils differs significantly between rat tail and bovine collagen, impacting MDA-MB-231 cells’ migration [23,24]. As of now, little is known about how the mechanical properties of the collagen matrix influence the chemotactic sensing of MDA-MB-231 cells to CXCL12 gradients. Furthermore, studies have shown cell shape can influence chemotaxis and taking into consideration MDA-MB-231 cells’ phenotypically heterogeneous nature, it remains to be understood how these morphological subpopulations respond to the chemokine gradient [25,26].

In this work, we investigate MDA-MB-231 cells response to several CXCL12 linear gradients (ranging from 0 to 500 ng/mL) embedded in type 1 bovine collagen matrices at two different concentrations, 1.5 and 2.0 mg/mL. Time-lapse images for 24 hours were captured and analyzed to obtain information on the dependence of cell shape on migration patterns [27].

## Materials and methods

### Materials

Human recombinant CXCL12 protein was purchased from Fujifilm Irvine Scientific. MDA-MB-231 cells were obtained from ATCC (catalog no. HTB-26™). The cell media, Dulbecco’s’ modified eagle medium (DMEM), was prepared in-house. Transwell chemotaxis migration assays were purchased from Millipore Sigma. Type I collagen (bovine, 5 mg/mL), 10x DMEM with phenol red, and chemotaxis μ-slides were purchased from Ibidi.

### Cell culture

MDA-MB-231 cells were maintained in DMEM supplemented with 10% FBS, 2 mM L-glutamine, and 1% penicillin-streptomycin. The cells were grown in 182 cm^2^ cell culture flask and the cell culture media was changed every 2-3 days. Cells were grown up to 70-90% confluency before they were cultured for chemotaxis assays.

### Transwell assay

The procedure adapted from the manufacturer’s protocol. Cells were resuspended in serum free DMEM to have a final concentration of 750,000 cells/mL. 100 µL of the cell solution was placed in the cell migration chamber plate and 150 µL of serum free media in the 96-well feeder tray. The cells were starved for 18-24 hours in the incubator at 37°C. The following day, the media in the feeder tray was replaced with 150 µL of chemoattractant media. After a 48-hour incubation at 37°C, the cell migration chamber plate was transferred to a new feeder tray with cell detachment buffer to displace the cells from the porous membrane. After 30 minutes, the transwell plate was removed from the incubator, lightly tilted back and forth to ensure the cells are completely detached. After, 50 µL of a 4X Lysis buffer/CyQuant GR Dye was added to the detachment buffer to lyse the cells and label their nucleic acids with a fluorescent dye. The mixture of each well was transferred to a plate suitable for fluorescence measurement and read on a plate reader at an excitation of 480 nm and emission of 520 nm.

### 3D chemotaxis assay

The procedure for creating the collagen matrix was adapted from Ibidi Application Note 26. Chemotaxis µ-slides were placed inside petri dishes with wet kimtech wipes to maintain humidity. Small aliquots of DMEM with and without FBS were placed in the incubator at 37°C overnight. The day of the experiment, the cells were washed twice with 1x phosphate buffer saline and detached from the flask using 0.25% EDTA-trypsin buffer. The cell pellet was resuspended in serum free DMEM (SF DMEM) at a final concentration of 12 million cells/mL. The reagents for the collagen gel mixture were placed on wet ice prior to mixing. The reagents were added based on the order of addition in Table 1, with excessive mixing in between each step to ensure homogenous distribution of collagen and cells. 6.5 µL of the collagen mixture was added to ports A and B (main channel) and plugged gently with tweezers. For gel polymerization, the slides were placed in the incubator for 1 hour at 37°C. After, 65 µL of SF DMEM was added to the reservoirs on both sides of the main channel and ng/mL of CXCL12 was added to the left reservoir in 15 µL increments to form a gradient. All experimental conditions were repeated at least three times. Cell movement was monitored on a Zeiss Axio Z1 Observer microscope with a 37 °C humidified heat chamber. Using bright-field microscopy, timelapse images were taken every 10 min for 24 hours.

**Table 1.**
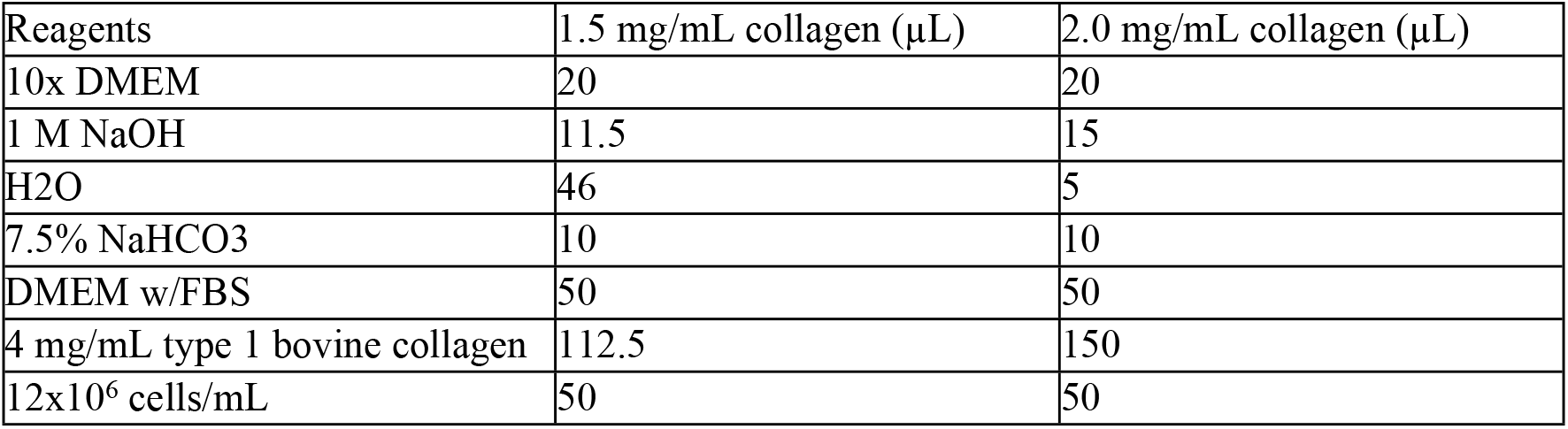
Order of addition for collagen gel mixture.

| Reagents | 1.5 mg/mL collagen (μL) | 2.0 mg/mL collagen (μL) |
| --- | --- | --- |
| 10x DMEM | 20 | 20 |
| 1 M NaOH | 11.5 | 15 |
| H<sub>2</sub>O | 46 | 5 |
| 7.5% NaHCO<sub>3</sub> | 10 | 10 |
| DMEM w/FBS | 50 | 50 |
| 4 mg/mL type 1 bovine collagen | 112.5 | 150 |
| 12x10<sup>6</sup> cells/mL | 50 | 50 |

### Data acquisition and statistical analysis

Timelapse images were analyzed using the automated tracking ImageJ software, CellTraxx.[27] Software version 4.8 is available on GitHub (https://github.com/borge-holme/celltraxx_download). CellTraxx pixel settings were adjusted for the Zeiss microscope. The 145-time frame videos were inverted for the cells to be identifiable with a dark outline prior to tracking. Cells that fit within the tracking parameters listed in Table 2 were included in the data analysis. At least 50 cells were tracked per replicate for each condition.

**Table 2.**
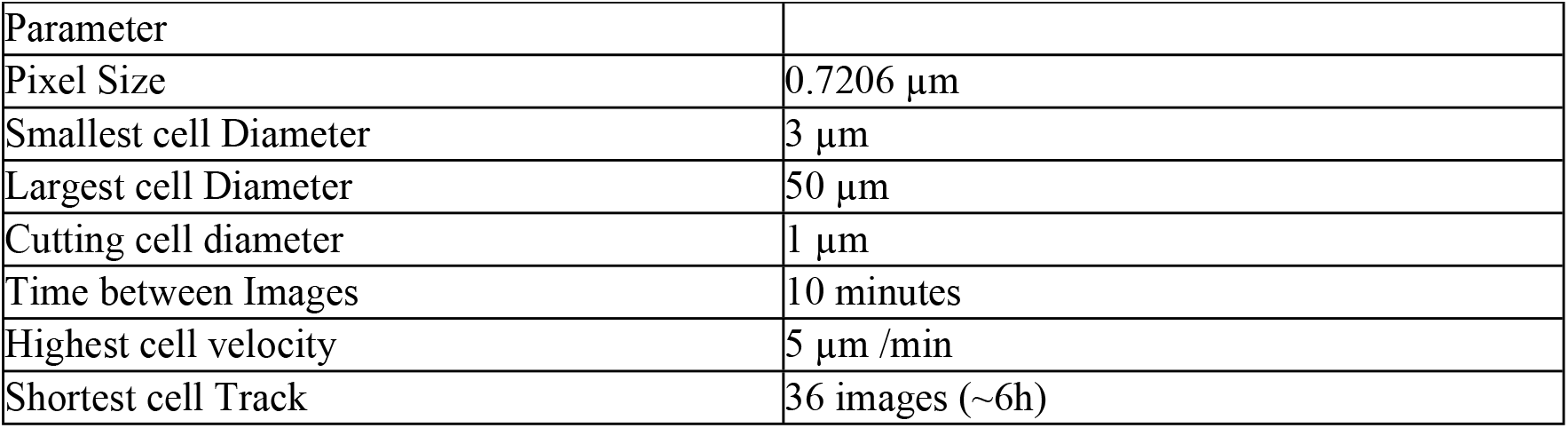
Significant parameters for single cell analysis using CellTraxx.

| Parameter |  |
| --- | --- |
| Pixel Size | 0.7206 $\mu\text{m}$ |
| Smallest cell Diameter | 3 $\mu\text{m}$ |
| Largest cell Diameter | 50 $\mu\text{m}$ |
| Cutting cell diameter | 1 $\mu\text{m}$ |
| Time between Images | 10 minutes |
| Highest cell velocity | 5 $\mu\text{m} / \text{min}$ |
| Shortest cell Track | 36 images (~6h) |

The forward migration index (FMI), which is the efficiency of the cell’s forward migration, and center of mass (COM), representing the average of single cell endpoints, were calculated to measure chemotaxis. Both FMI and COM were calculated parallel to the gradient, (x-direction), and perpendicular to the gradient (y-direction) where n is the total number of individual cells, x_i,end_ and y_i,end_ are the cell’s endpoint coordinates, and d_i,accum_ is the total distance the cell has traveled [28].

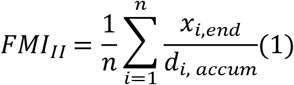

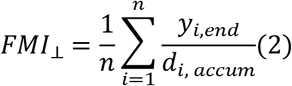

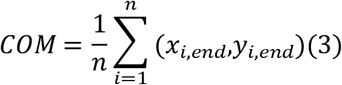

Significant chemotactic activity was determined if (1) FMI_II_ was greater than the FMI_⊥_and (2) FMI value of the CXCL12 gradient was larger than the negative control. Cell velocity was measured to monitor changes in cell movement. For statistical analysis, an ordinary one-way analysis of variance (ANOVA) test was performed followed by a Dunnett’s multiple comparison test. Figure plots and statistical analysis was plotted using GraphPad Prism 9.

## Results and Discussion

### CXCL12 promotes chemotactic activity of TNBC cells in a 2D migration assay

An initial assessment of the chemotactic effects of CXCL12 gradients on MDA-MB-231 cells was conducted using a transwell assay with a fluorometric readout for quantitative analysis (Fig 1A). Several concentrations were tested, ranging from 0 to 500 ng/mL, over a 48-hour time period in a serum free environment to enhance the cells’ sensitivity to the chemokine gradient. In Fig 1B, the number of cells migrating into the lower chamber increased as the concentration of CXCL12 increased, with a significant migration occurring between 200-500 ng/mL and the highest chemotactic activity occurring at 200 ng/mL. This optimal concentration range is similar to previously reported results in other 2D migration assays [29]. There are significant drawbacks of utilizing this assay, particularly, the inability to measure the cell’s motility in real time and the formation of a steep gradient, which is not comparable to the gradients formed *in vivo*.

**Figure 1.**
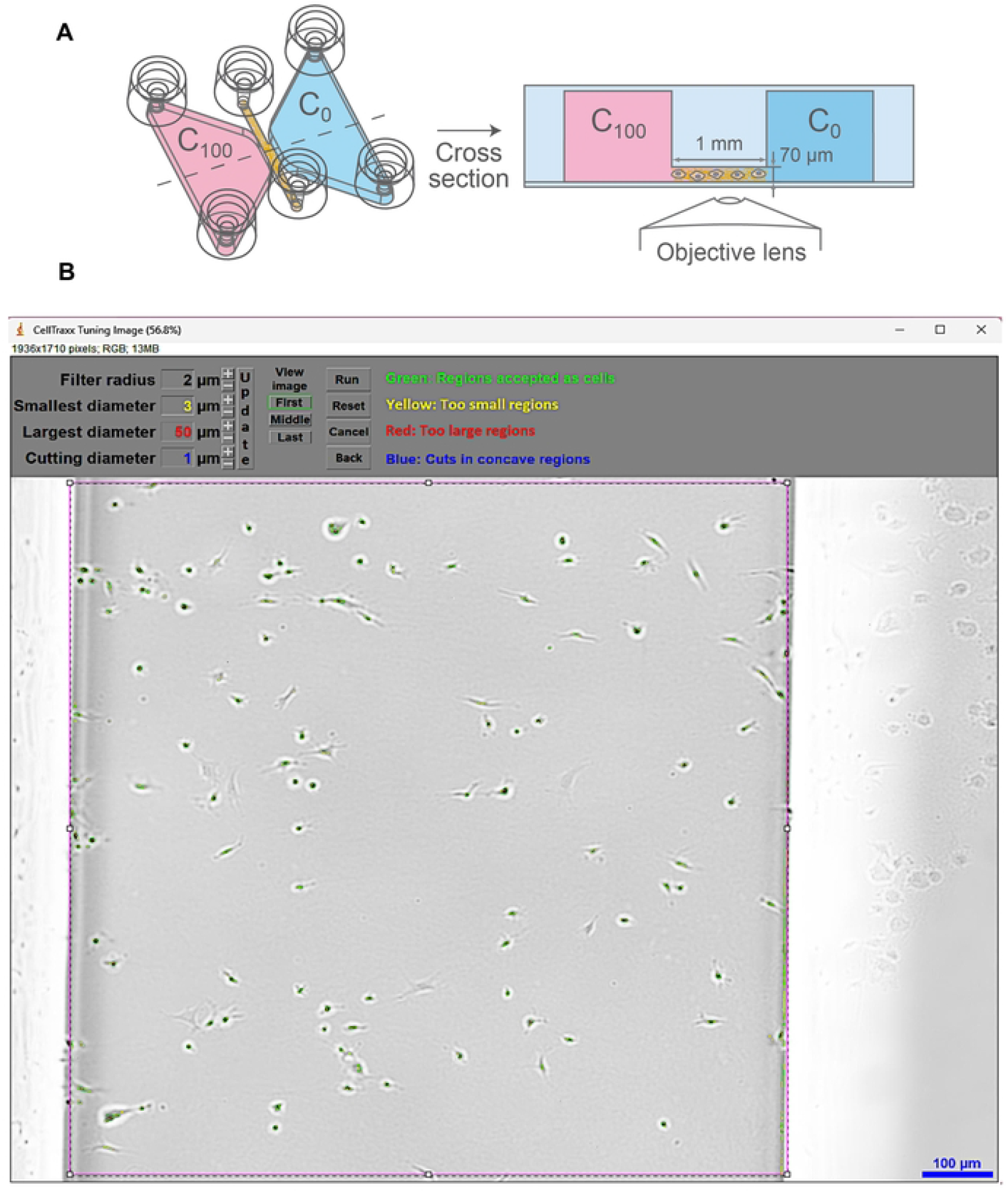
2D migration setup and migration analysis. (A) Schematic of 2D transwell migration assay made in BioRender and (B) chemotactic response of MDA-MB-231 cells to seven different CXCL12 gradients ranging from 0 to 500 ng/mL. 7.5×10^4^ cells/well migrated towards the lower chamber for 48 hours. Fluorescence measurements were normalized to the negative control. Error bars are represented as standard deviation (n=3).*p<0.05,**p<0.01, Dunnett’s multiple comparison test.

Therefore, the concentrations of CXCL12 yielding the most cell migration in this 2D assay were applied to a 3D cell culture system as linearly diffused gradients.

### Tracking individual cells in a 3D matrix using CellTraxx

3D migration was observed using the ibidi µ-slide for chemotaxis (Fig 2A). This device establishes a long-term linear gradient that can diffuse through a 3D matrix and monitor individual cells’ migration and morphology over time [30]. Type 1 bovine collagen was used as the 3D matrix to mimic the ECM. Studies have shown collagen matrices from the same source material at different concentrations can impact breast cancer cell invasion [23]. Therefore, we tested collagen concentrations, 1.5 mg/mL and 2.0 mg/mL, to determine how collagen density could impact cell behavior in the presence of a chemoattractant gradient. MDA-MB-231 cells are known for their cellular heterogeneity and dynamic movement, making it difficult and time-consuming to monitor these cells using manual tracking. Thus, to accurately identify and track these cells for 24 hours, the automated software CellTraxx was used as a macro plugin in ImageJ. Time-lapse images were uploaded to CellTraxx version 4.8 to measure the cell’s aspect ratio in addition to other migration parameters (Fig 2B). Extensive optimization was conducted for the CellTraxx settings “highest cell velocity” and “time between images” to ensure an accurate track of an individual cell’s chemotactic movement.

**Figure 2.**
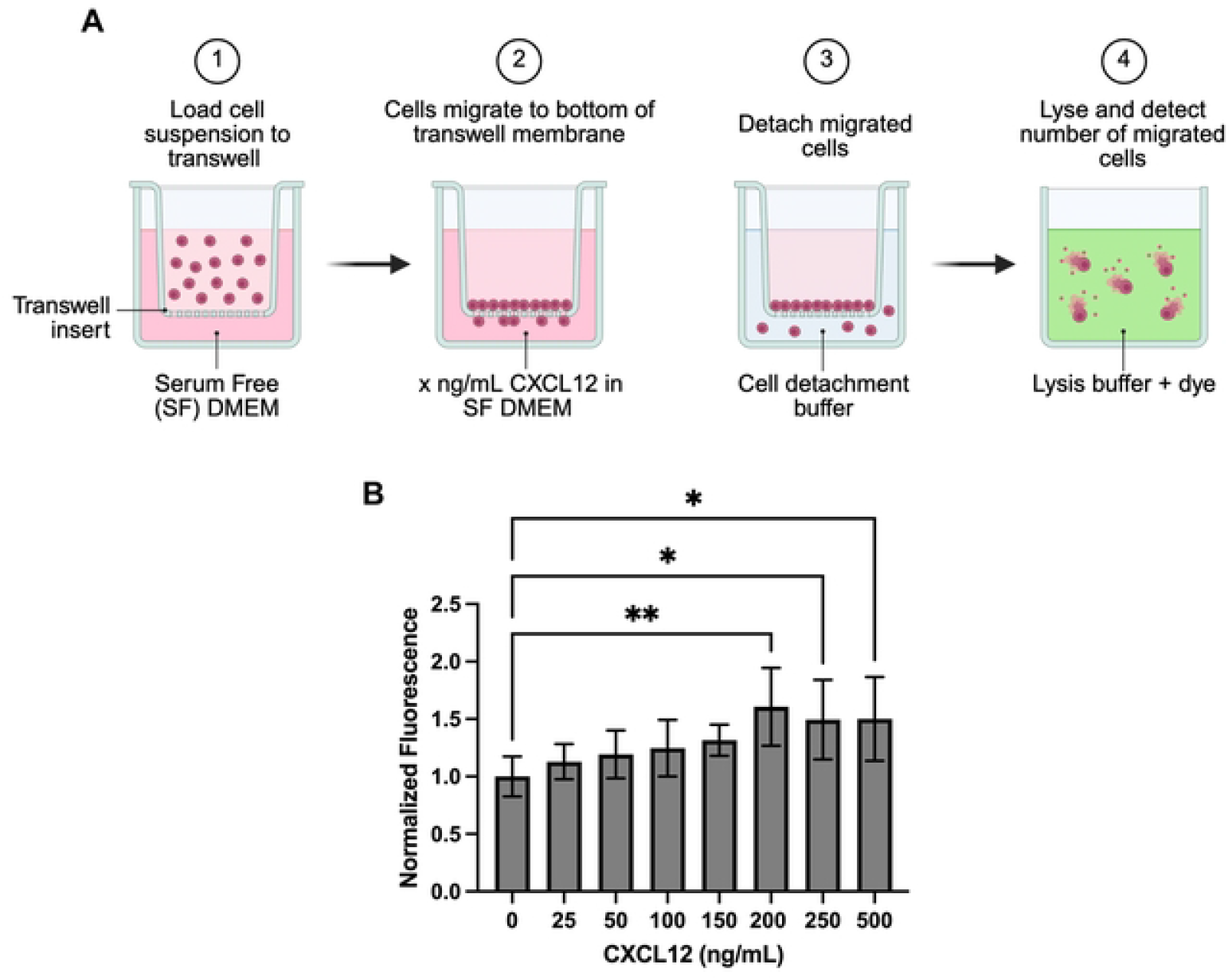
3D migration setup and cell tracking. (A) Diagram of the ibidi µ-slide for chemotaxis (Image courtesy of ibidi GmbH) (B) Snapshot of CellTraxx tuning interface. Zeiss microscope images were inverted to make the cells outline detectable by the CellTraxx software.The formatted video was uploaded to CellTraxx version 4.8. The CellTraxx settings were adjusted to accurately identify MDA-MB-231 cells shape and movement.

Initial cell tracking was conducted on the entire cell population in the 1.5 and 2.0 mg/mL collagen concentrations. Globally, for the cells embedded in the 1.5 mg/mL collagen gel in the presence of no chemoattractant, the FMI_II_ is 0.001 ± 0.014 and FMI_⊥_ is -0.006±0.004 (Fig 3A). The COM_II_ and COM_⊥_ are -0.508± 1.774 µm and -0.366± 3.179 µm, respectively as shown in Fig 3B. Surprisingly, small amounts of CXCL12, about 50 ng/mL, induced a strong FMI_II_ in the negative direction with a value of -0.020±0.005. The COM_II_ and COM_⊥_ are -5.018±0.947 µm and -0.045±0.509 µm, respectively (Figs 3A and 3B). As the concentration of CXCL12 increased beyond 50 ng/mL, FMI_II_ progressively increased in a dose-dependent manner, while FMI_⊥_ correspondingly decreased. The chemotactic response reaches a maximal level at approximately 300 ng/mL CXCL12; the FMI_II_ value of 0.015± 0.005 and FMI_⊥_ of -0.006± 0.002, fits the first criteria of chemotaxis. The COM_II_ and COM_⊥_ are 1.668± 0.638 µm and -0.159± 0.581µm, respectively. We note that the FMI values are quite small, and it is definitely peculiar that the cells moved backwards at 50 ng/mL CXCL12 in this assay, which might suggest that at this concentration, CXCL12 can act as a chemorepellent in low-density matrices; it is also peculiar that the perpendicular values for FMI and COM were not always zero. This latter result suggests that the chemical gradient may not have been strictly formed in the x-direction, in spite of our many efforts to ensure this [31].

**Figure 3.**
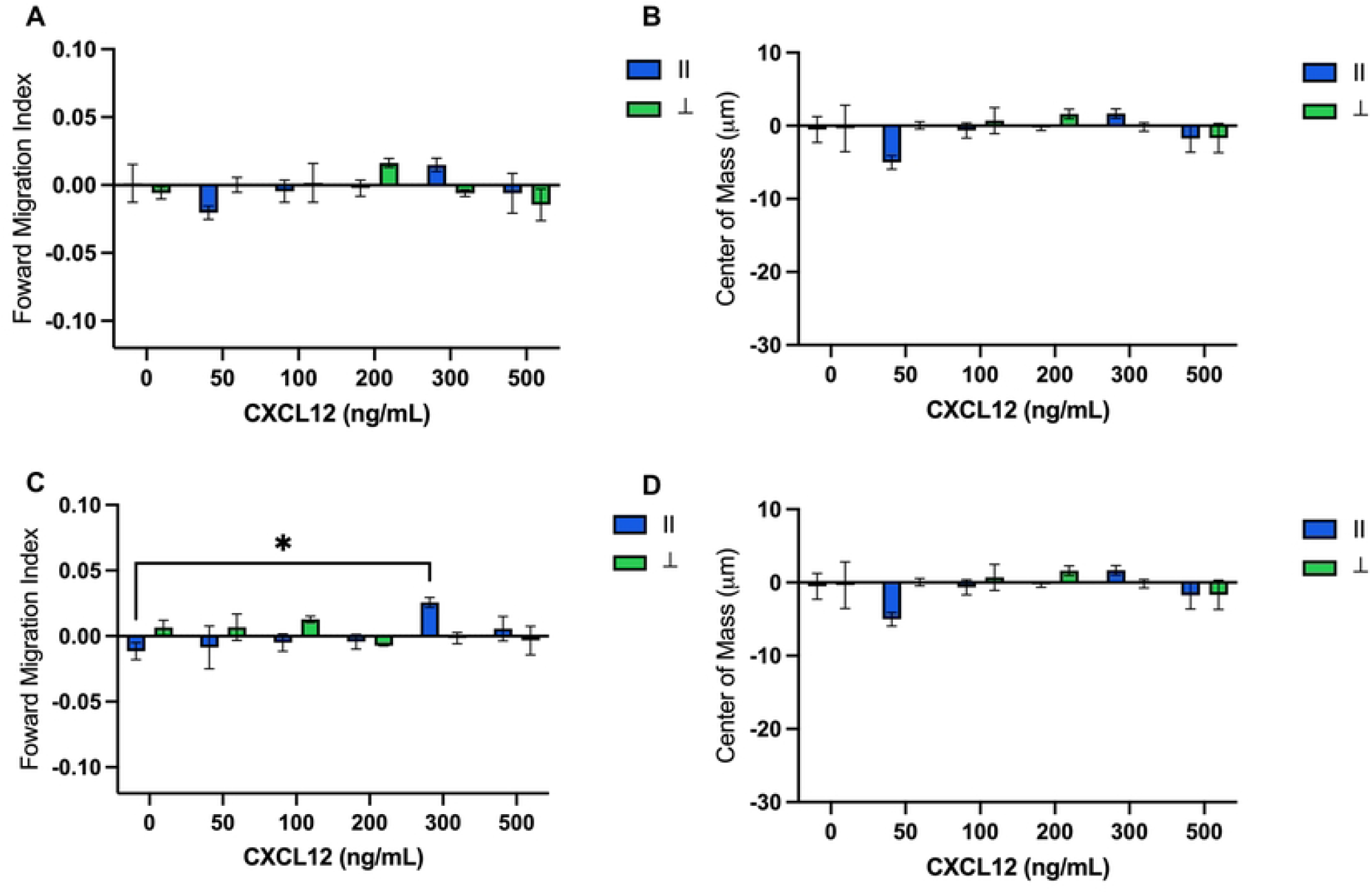
FMI and COM of the entire cell population exposed to several CXCL12 gradients. (A) Forward Migration Index and (B) center of mass of the entire MDA-MB-231 cell population in the presence of a CXCL12 linear gradients in 1.5 mg/mL collagen gel. (C) and (D) FMI and COM in 2.0 mg/mL conditions. Blue bars represent FMI and COM parallel (∥) to the gradient and green bars represent the perpendicular (⍰) to the gradient. *p<0.05, Dunnett’s test. Error bars represented as error of mean.

For 2.0 mg/ml collagen concentration conditions, the FMI_II_ is -0.011± 0.007 and FMI_⊥_ is -0.007± 0.006 when there is no chemoattractant (Fig 3C). The COM_II_ and COM_⊥_ are -1.304± 0.652 µm and .0707± 0.986 µm, respectively (Fig 3D). The FMI_II_ and COM_II_ has a relatively more negative value in comparison to the 1.5 mg/mL collagen negative control, which can be attributed to the increase in collagen density. However, the trends are similar in both conditions, where the FMI_II_ gradually increases as the FMI_⊥_ decreases. It was observed 300 ng/mL significantly promoted the forward migration of the TNBC cells with a FMI_II_ value of 0.026±0.004 and FMI_⊥_ is -0.007±0.006. The COM_II_ and COM_⊥_are 3.013± 0.967µm and -1.014± 1.231µm, accordingly. A tighter standard deviation was noticed across all conditions, in comparison to the 1.5 mg/mL collagen matrix. The apparent “backwards” cell movement at ∼50 ng/mL CXCL12 for the 1.5 mg/mL collagen matrix was not evident at 2.0 mg/mL collagen concentration, suggesting that apparent chemorepellent behavior of a nominal chemoattractant is ECM-dependent. Overall, cell migration of these triple-negative breast cancer cells was rather modest, with only 300 ng/mL CXCL12 at a collagen density of 2.0 mg/mL showing statistical significance.

### MDA-MB-231 cells response to CXCL12 is shape dependent

One of the initial steps of TNBC metastasis is the epithelial-to-mesenchymal transition (EMT), during which cells undergo extensive cytoskeletal reorganization from a rounded epithelial shape to an elongated mesenchymal shape [32]. This process enables the cells to detach from the primary tumor, intravasate into the vasculature, and proliferate at a secondary site [33]. It is well known that the CXCL12/CXCR4 axis induces EMT of breast cancer cells [34,35]. However, it is yet to be understood how the cell morphologies within the heterogeneous population respond to the chemoattractant gradient.

Based on the initial assessment of the entire cell population, we hypothesize that the cell’s morphology may be contributing to the large deviations of migration behavior found across the tested gradients. To test this hypothesis, the average aspect ratio (AR) for each cell was calculated, plotted on histogram plot and assigned a cell shape: round, ellipsoidal, and elongated (Fig 4A). The median of the histogram plot, AR between 3.5 and 6.0, was assigned as the ellipsoidal due to the cell’s constant retraction and contraction behavior. Cell with an AR less than 3.5 was classified as round, where the cells remain in the round shape throughout the duration of the experiment. Cells with AR more than 6.0 were classified as elongated, due to the cell’s elongation shape and consistent deformation of the nucleus as it traverses the collagen. As shown in Fig 4B, there is a difference in the cell shape population within the two different collagen matrices for each gradient. With no chemoattractant present, the percentage of round cells remained similar for both collagen matrices, a bulk of cells were ellipsoidal at the 2.0 mg/mL, and the majority were elongated in the 1.5 mg/mL collagen gel. Although the percentages for the ellipsoidal cell shape remained similar for each gradient, there was a noticeable difference for the round and elongated cells when exposed to the 300 ng/mL CXCL2 gradient, where there was increase in the round cells at 2 mg/mL collagen and a decrease in the 1.5 mg/mL collagen conditions (Table S1). After classification of cell shapes, the chemotaxis data was subsequently re-analyzed, and averages were calculated for each group. Since the gradient was established in the x-direction, only the FMI_II_ and COM_II_ are highlighted in this section for the different cell shapes (denoted as FMIx and COMx). Cell velocity was also included in our measurements to determine if cell migration speed differs between cell shape.

**Figure 4.**
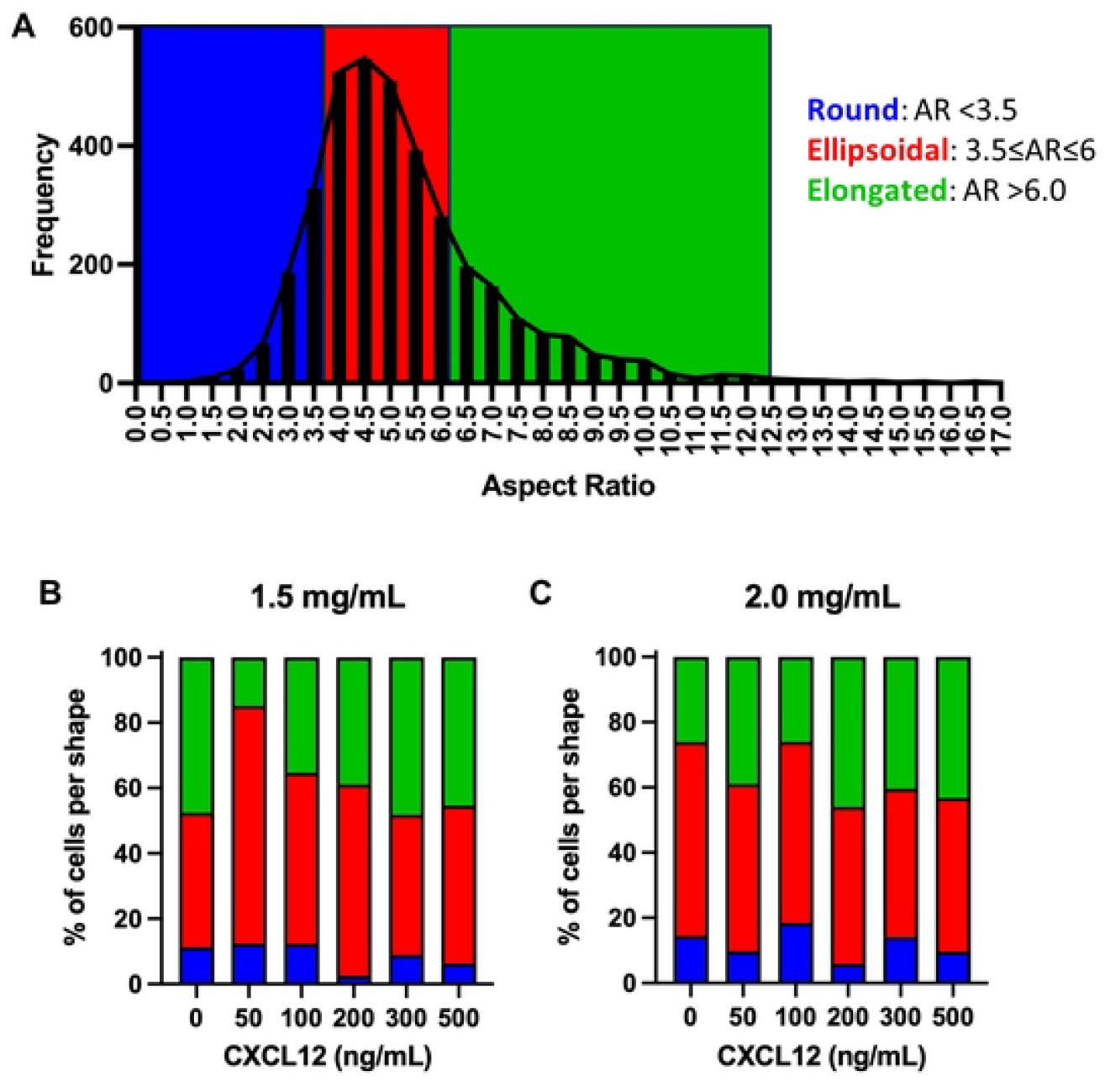
Distribution of different cell shapes. (A) Histogram plot of the aspect ratio (AR) of all single cells. Cell shape was divided into three AR ranges: round (AR less than 3.5), ellipsoidal (AR between 3.5 and 6.0), and elongated (AR more than 6.0). (B) Percentage of each cell shape population for 1.5 mg/mL and (C) 2.0 mg/mL type 1 bovine collagen conditions derived from the histogram. Blue bars represent round cells, red bars represent ellipsoidal cells, and green bars represent elongated cells.

### Migratory behavior of round cells

The round cells display few protrusions in their cytoskeleton, leading to limited ECM-cell adhesions. The cells behaved similarly in both collagen matrices across the CXCL12 gradients but responded differently when they were exposed to increasing amounts of CXCL12. As shown in Fig 5A, there are fluctuations in the FMI as the cells experience CXCL12 concentrations of ≥100 ng/mL, where there is a more consistent upward trend in the 2.0 mg/mL collagen conditions. In comparison to the ellipsoidal and mesenchymal cells, the round cells have a relatively higher COM value when exposed to high CXCL12 concentrations, suggesting that the cells spend less time interacting with ECM (Fig 5B). Interestingly, the velocity decreases when the round cells are exposed to CXCL12 gradients >200 ng/mL and migrate significantly slower in comparison to when there is no chemoattractant present. Additionally, the averaged velocities of the round cells are less than the reported values of the ellipsoidal and elongated cells, which contradicts previous works that suggest cells exhibiting an ameboid migration move relatively faster [19].

**Figure 5.**
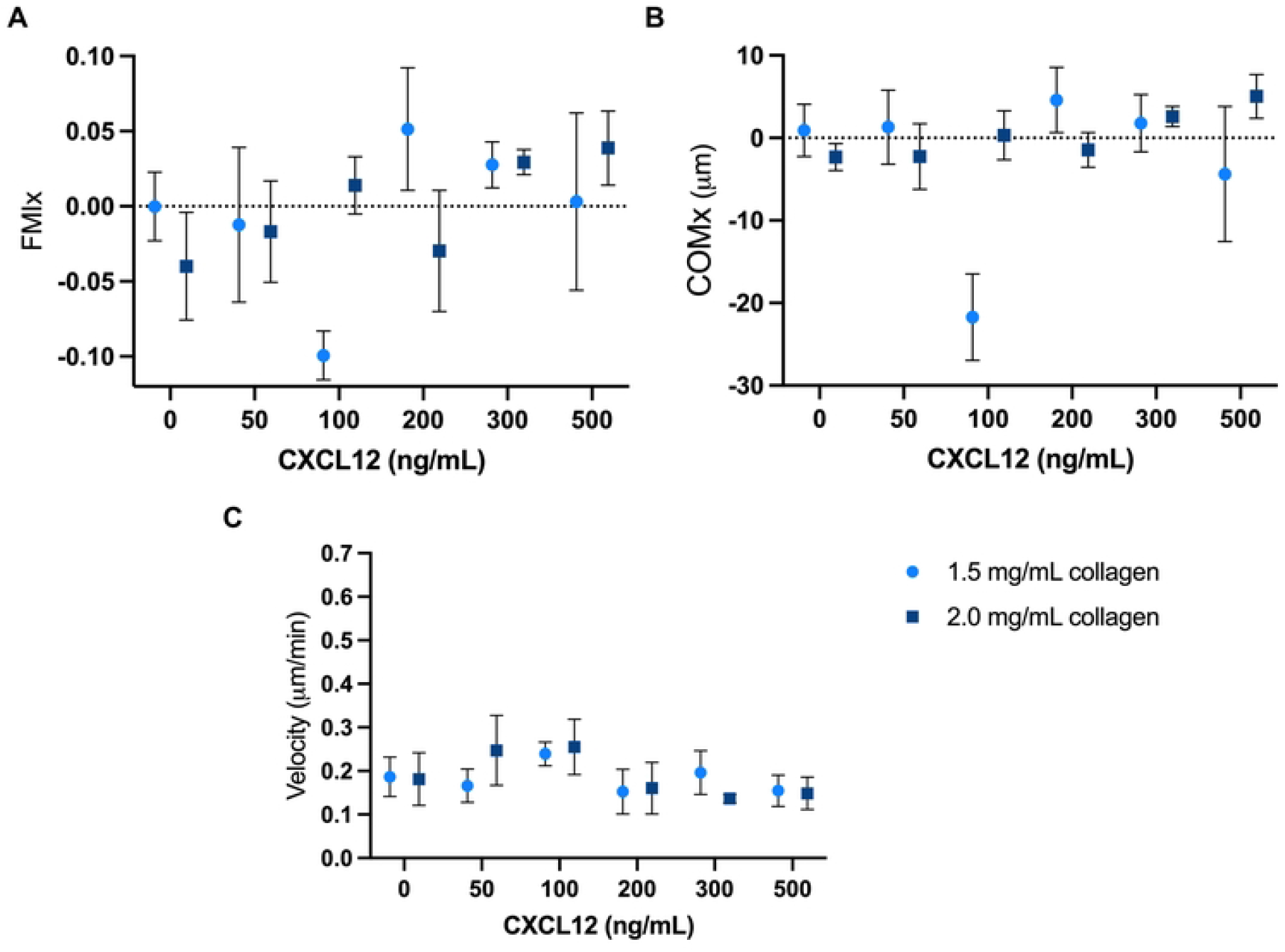
Migration analysis of round cells. (A) FMI, (B) COM, and (C) Velocity of the round shaped MDA-MB-231 cells in the presence of a CXCL12 linear gradients. Error bars represent standard error of mean.

### Migratory behavior of ellipsoidal cells

Ellipsoidal cells are denoted as cells that exhibit both round and elongated features, where there is partial elongation, but do not display a deformed nucleus. This group demonstrated the most chemokinetic activity, where the cells are almost non-responsive to any of the CXCL12 gradients in both collagen matrices, as shown in the FMI (Fig 6A). Rather than moving across the collagen reservoir, these ellipsoidal cells typically remained in place and migrated in a circular motion, often returning to their originating spot, hence why there is little to no change in the COM (Fig 6B). However, in this constant circular motion, we observe these cells display higher velocities than the round cells. Additionally, we noticed the velocity of this subpopulation behave similarly to the round cells when a gradient of ≥200 ng/mL was introduced (Fig 6C). In the 1.5 mg/mL matrix, the average velocities are consistent across the biological replicates from 100 to 300 ng/mL. Interestingly, there is a switch in migratory behavior where the ellipsoidal cells in the denser collagen matrix migrate slightly faster towards the high CXCL12 gradients.

**Figure 6.**
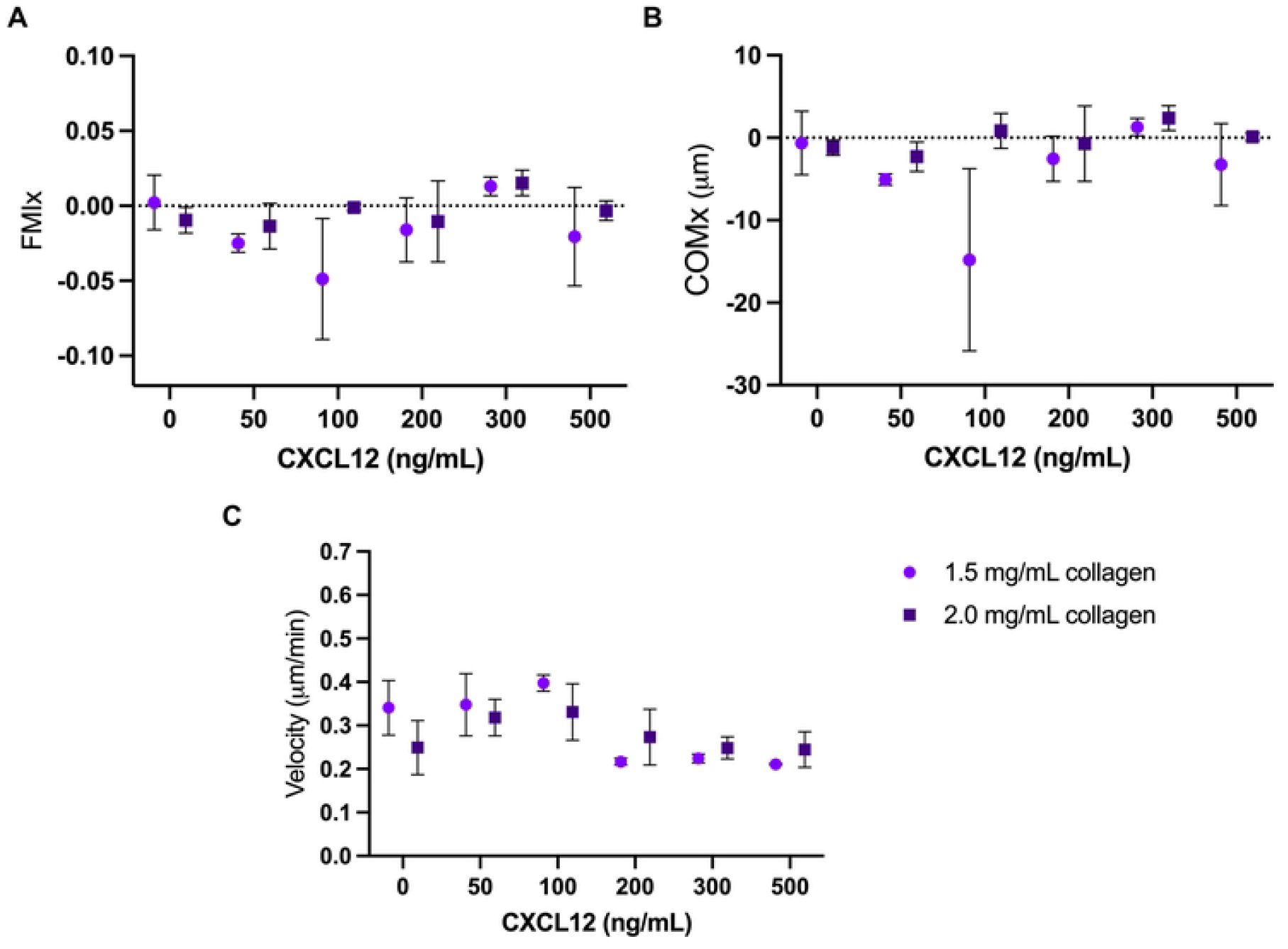
Migration analysis of ellipsoidal cells. (A) FMI, (B) COM, and (C) Velocity of the ellipsoidal shape MDA-MB-231 cells in the presence of a CXCL12 linear gradients. Error bars represent standard error of mean.

### Migratory behavior of elongated cells

Elongated cells display a deformable nucleus and reorganize their cytoskeleton to have a spindle-like shape. Between 0-50 ng/mL, the cells have large fluctuations in their FMI in the 1.5 mg/mL collagen matrix than the 2.0 mg/mL collagen matrix. The elongated cells seem to favor the denser matrix with a 300 ng/mL chemokine gradient as denoted by the FMI value of 0.037, which is the largest value observed across all the conditions (Fig 7A). As the averaged distance traveled of the elongated cells increased towards the high CXCL12 gradients, similar COM values were measured at CXCL12 concentrations of 300 ng/mL in both collagen matrices. As the CXCL12 concentration increases, we observed smaller standard deviations across the biological replicates, indicating this is a consistent migratory behavior for the cells (Fig 7B). When the cells interacted with the low CXCL12 gradients, between 0-100 ng/mL, there was a noticeable difference in cell velocities, where the cells migrate faster in the 1.5 mg/mL than in the 2.0 mg/mL collagen gel (Fig 7C). For the high CXCL12 gradients, between 300-500 ng/mL, the reported average velocities are identical in both matrices, calculated to be in a range of 0.2 to 0.3 µm/min. In the denser matrix, the cell velocities remain the same across all gradients, suggesting the cells are strongly interacting with the ECM while simultaneously sensing the chemoattractant.

**Figure 7.**
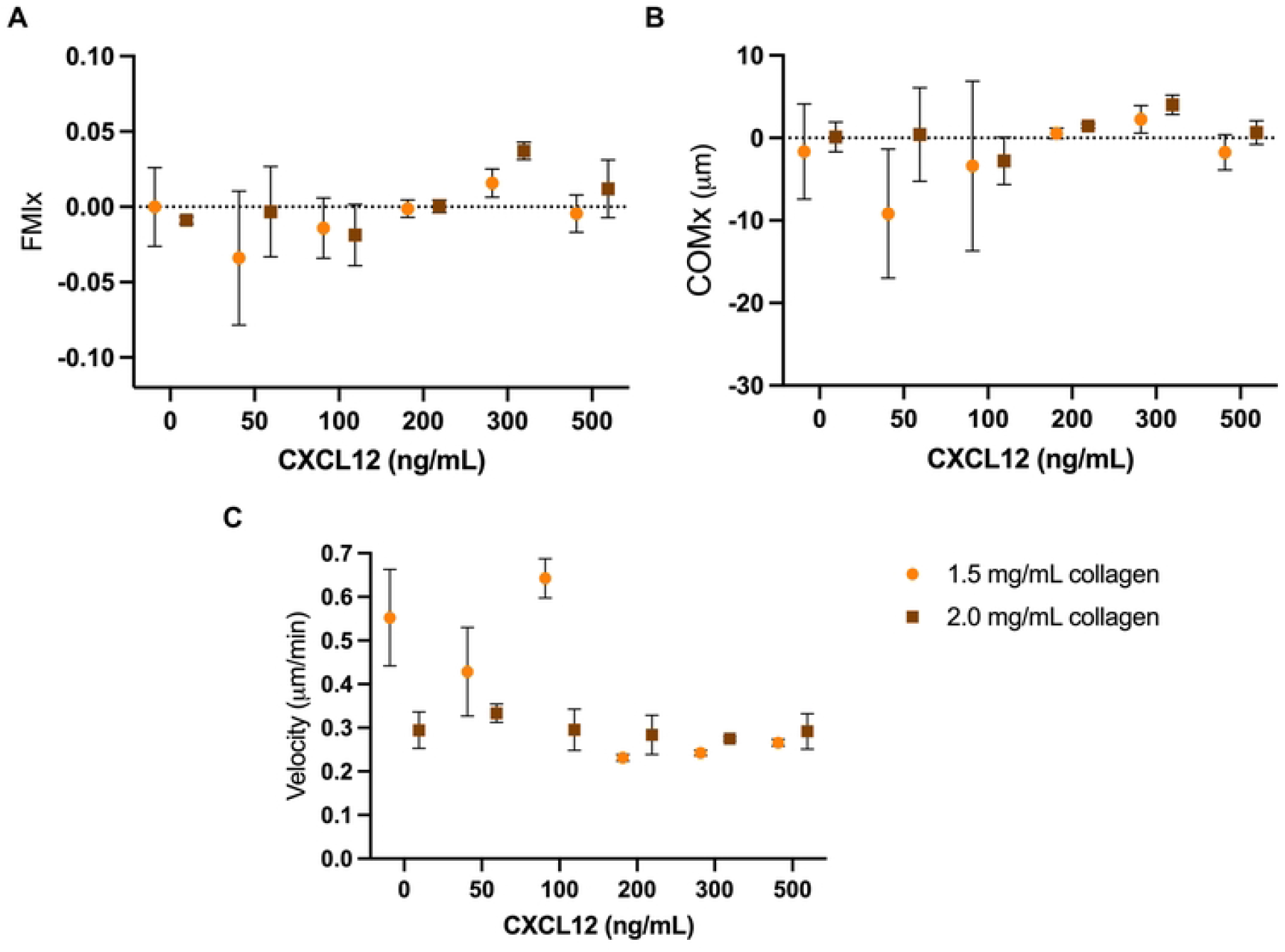
Migration analysis of elongated cells. (A) FMI, (B) COM and (C) Velocity of the elongated shape MDA-MB-231 cells in the presence of a CXCL12 linear gradients. Error bars represent standard error of mean.

## Conclusion

In this work, we explored the relationship between cell shape and chemotactic cell migration in a type 1 bovine collagen matrix. We observed CXCL12 by itself is not a potent chemoattractant in a 3D system in comparison to a 2D system, supporting the idea that the mechanism for 2D migration cannot be directly translatable to a 3D environment. In both collagen gel concentrations, the cells exhibited an overall random migration when exposed to CXCL12 gradients in a dose-dependent manner. Nevertheless, the entire cell population preferred to migrate forward in the denser 2.0 mg/ml collagen gel. Interestingly, when the cells were categorized based on their shape, we found the round and ellipsoidal cells did not have a change in the average FMIx in the denser matrix but the average FMIx of the elongated cells increase over two-fold in the denser matrix when exposed to a 300 ng/mL CXCL12 gradient.. Our findings suggest there is a need to explore the relationship between the chemotaxis of heterogeneous TNBC cells and metastasis using complex 3D ECM mimics. A potential future direction would be to incorporate cells that generate CXCL12 gradients in the collagen gel to monitor real-time gradient formation in a biologically diverse environment.

## Acknowledgments

We thank Dr. Sandy McMasters from the UIUC Cell Media Facility for preparing cell culture materials and helpful discussion.

## Supporting Information

**S1 Table. Tabulated results of migratory analyses for 1.5 mg/mL and 2.0 mg/mL collagen matrix conditions**.

**S1 Fig. Representative trajectory plots of MDA-MB-231 cells embedded in 1.5 mg/mL collagen matrix**. (A) 0 ng/mL (B) 50 ng/mL (C) 100 ng/mL (D) 200 ng/mL (E) 300 ng/mL (F) 500 ng/mL CXCL12 gradients.

**S2 Fig. Representative trajectory plots of MDA-MB-231 cells embedded in 2.0 mg/mL collagen matrix**. (A) 0 ng/mL (B) 50 ng/mL (C) 100 ng/mL (D) 200 ng/mL (E) 300 ng/mL (F) 500 ng/mL CXCL12 gradients.

**S1 Movie. Time-lapse video of MDA-MB-231 cells exposed to the negative control in 2.0 mg/mL with CellTraxx**.

**S2 Movie. Time-lapse video of MDA-MB-231 cells exposed to the 300 ng/mL CXCL12 gradient in 2.0 mg/mL with CellTraxx**.

